# Ankyrin2 is required for neuronal morphogenesis and long-term memory and interacts genetically with HDAC4

**DOI:** 10.1101/2021.07.18.452850

**Authors:** Silvia Schwartz, Sarah J Wilson, Tracy K Hale, Helen L Fitzsimons

## Abstract

Dysregulation of *HDAC4* expression and/or subcellular distribution results in impaired neuronal morphogenesis and long-term memory in *Drosophila melanogaster*. A recent genetic screen for genes that interact in the same molecular pathway as *HDAC4* identified the cytoskeletal adapter *Ankyrin2* (*Ank2*). Here we sought to investigate the role of *Ank2* in neuronal morphogenesis, learning and memory, and to examine the nature of interaction with *HDAC4*. We found that Ank2 is expressed widely throughout the *Drosophila* brain where it localizes predominantly to axon tracts. Pan-neuronal knockdown of *Ank2* in the mushroom body, a region critical for memory formation, resulted in defects in axon morphogenesis, and similarly reduction of *Ank2* in lobular plate tangential neurons of the optic lobe disrupted dendritic branching and arborization. Conditional knockdown of *Ank2* in the mushroom body of adult *Drosophila* significantly impaired long-term courtship memory, and this requirement for *Ank2* was isolated to gamma (γ) neurons of the mushroom body. As overexpression of *HDAC4* in γ neurons also impairs the formation of long-term courtship memory, this suggests that any functional relationship between these proteins during LTM likely occurs in γ neurons. We determined that the genetic interaction requires the presence of nuclear *HDAC4* and is not dependent on a conserved putative ankyrin-binding motif present in HDAC4. In summary, we provide the first characterization of the expression pattern of Ank2 in the adult *Drosophila* brain and demonstrate that Ank2 is critical for morphogenesis of the mushroom body and for the molecular processes required in the adult brain for formation of long-term memories.

## Introduction

Histone deacetylase 4 (HDAC4) is a member of the Class IIa family of histone deacetylases, which are categorized by the presence of an extended N-terminal regulatory region and their ability to undergo nucleocytoplasmic shuttling (Grozinger and Schreiber 2000; McKinsey *et al*. 2000; Wang *et al*. 2000; Wang and Yang 2001; Chawla *et al*. 2003; Schlumm *et al*. 2013). HDAC4 is expressed widely throughout the brain and dysregulation of HDAC4 expression and/or subcellular distribution has been implicated in several neurodevelopmental and neurodegenerative disorders (Williams *et al*. 2010b; Li *et al*. 2012; Sando *et al*. 2012; Herrup *et al*. 2013; Mielcarek *et al*. 2013; Shen *et al*. 2016; Trazzi *et al*. 2016). Loss of *HDAC4* is associated with Chromosome 2q37 deletion syndrome [MIM 600430], the clinical features of which include developmental delay, autistic features and intellectual disability (Williams *et al*. 2010b; Williams *et al*. 2010a; Morris *et al*. 2012a; Morris *et al*. 2012b; Villavicencio-Lorini *et al*. 2013). Developmental delay is the most common feature, and while haploinsufficiency of *HDAC4* displays variable penetrance with respect to intellectual disability (Wheeler *et al*. 2014), the predominant genetic cause is considered to be loss of function of *HDAC4* (Dag *et al*. 2019). HDAC4 has been shown to regulate synaptic plasticity and memory formation in several animal models (Wang *et al*. 2011; Kim *et al*. 2012a; Sando *et al*. 2012; Fitzsimons *et al*. 2013; Schwartz *et al*. 2016; Main 2021); In mice, brain-specific conditional knockout of *HDAC4* results in impaired spatial memory (Kim *et al*. 2012b; Kim *et al*. 2012a) and in *Drosophila*, RNAi knockdown of *HDAC4* in the brain also impairs long-term memory (LTM) formation (Fitzsimons *et al*. 2013). Disruption of normal subcellular distribution is also detrimental to neurodevelopment and cognition; a recent study identified seven individuals presenting with developmental delay, intellectual disability and other neurodevelopmental deficits, all of whom had *de novo* mutations in the 14-3-3 binding site within HDAC4. This motif is required for phosphorylation-dependent nucleocytoplasmic shuttling of HDAC4, and thus the disruption to 14-3-3 binding is presumed to disrupt normal HDAC4 shuttling, leading to nuclear accumulation (Wakeling *et al*. 2021). In mice, a truncated mutant of HDAC4 that accumulates in the nucleus causes deficits in spatial memory. This mutant lacks the deacetylase domain (Sando *et al*. 2012), however vertebrate HDAC4 is catalytically inactive (Lahm *et al*. 2007; Bottomley *et al*. 2008), reviewed by (Fitzsimons 2015). Similarly, overexpression of *HDAC4* in the adult mushroom body, a brain region essential for memory formation in *Drosophila* (Heisenberg *et al*. 1985; McBride *et al*. 1999), prevented LTM formation, as did a catalytically inactive mutant (Fitzsimons *et al*. 2013). This memory phenotype was recapitulated with a mutant of *HDAC4* that is restricted to the nucleus, and when expressed during development it also impaired morphogenesis of mushroom body axons (Main 2021).

Increased nuclear HDAC4 has also been observed in hippocampal pyramidal neurons in mouse models of Alzheimer’s disease (Shen *et al*. 2016). Moreover, HDAC4 was found to accumulate in pyramidal cells and layer III of the frontal cortex in post-mortem brains from individuals with Alzheimer’s disease, with the abundance of nuclear HDAC4 correlating with the clinical dementia scores (Herrup *et al*. 2013; Shen *et al*. 2016). These data together indicate that nuclear HDAC4 impairs cognitive function and that dysregulation of nucleocytoplasmic shuttling contributes to disease progression.

To gain further knowledge of the molecular pathway through which HDAC4 acts, we previously carried out a genetic enhancer screen in *Drosophila* photoreceptors for genes that interact genetically with *HDAC4* (Schwartz *et al*. 2016). Expression of *HDAC4* in the eye results in a mild “rough” eye phenotype, as characterized by disruption of the regular ommatidial patterning. We identified a group of cytoskeletal regulators including *Ankyrin2* (*Ank2*), that when knocked down in combination with *HDAC4* overexpression resulted in an enhanced rough eye phenotype, indicative of a genetic interaction. We focused our attention on further investigating *Ank2*, since the altered expression of the human homologue *ANK3* has been associated with a variety of neurodevelopmental disorders including intellectual disability, epilepsy, attention deficit hyperactivity disorder (Iqbal *et al*. 2013), bipolar disorder (Tesli *et al*. 2011) and autism spectrum disorder (Bi *et al*. 2012). Moreover, *ANK3* single nucleotide polymorphisms have been associated with schizophrenia (Athanasiu *et al*. 2010; Yuan *et al*. 2012; Nie *et al*. 2015a; Guo *et al*. 2016; Hughes *et al*. 2018) and Alzheimer’s disease (Morgan *et al*. 2008).

Ankyrins are adapter proteins that link the underlying spectrin-actin cytoskeleton to integral membrane proteins such as ion channels, anion exchangers, signaling proteins and cell adhesion molecules (Bennett 1978; Bennett and Baines 2001; Mohler *et al*. 2002). Canonical ankyrins are comprised of an N-terminal membrane binding domain, which contains 24 ANK repeat motifs organized as two anti-parallel α-helices that mediate protein-protein interactions. They also contain a central spectrin-binding domain, a death domain and a less conserved C-terminal regulatory domain (Sedgwick and Smerdon 1999; Cunha and Mohler 2009). *Drosophila Ank2* displays high similarity to human *ANK3* (Iqbal *et al*. 2013), sharing 57% amino acid identity over the whole protein and 71.2% identity across the ankyrin repeat region. Similarly to *ANK3*, expression of *Ank2* is restricted to neurons and a number of transcript isoforms are also expressed. In the embryonic nervous system, shorter isoforms of *Ank2* (Ank2-S) localize to the cell bodies of neurons, whereas medium Ank2-M isoforms localize to axons and are essential for viability (Hortsch *et al*. 2002). A longer Ank2-L isoform containing an additional C-terminal domain localizes to axons and synaptic boutons at the neuromuscular junction. Mutants with disrupted Ank2-L expression display a loss of synapse stability as well as a reduction in the terminal bouton size, disassembly of presynaptic active zone and retraction of the synaptic microtubule cytoskeleton (Koch *et al*. 2008; Pielage *et al*. 2008), thus Ank2-L is required for synapse stability and normal morphology at the neuromuscular junction in *Drosophila* larvae.

Despite comprehensive investigation of the roles that Ank2 plays in neuronal development, the role of Ank2 in the adult brain and functional consequences of reduced Ank2 has received little attention. Here we aimed to investigate the role of Ank2 in learning, memory and development of the *Drosophila* brain as well as to examine the nature of the interaction between HDAC4 and Ank2 and to determine whether this is an important mechanism through which HDAC4 regulates neuronal development and memory.

## METHODS

### Fly strains

All flies were raised on standard medium on a 12-hour light/dark cycle and maintained at a temperature of 25°C unless otherwise indicated. *w[*]; P{w[+mW.hs]=GawB}OK107 eγ[OK107]/In(4)ci[D], ci[D] pan[ciD] sv[spa-pol] (OK107-GAL4), w[*]; P{w[+mC]=GAL4-ninaE.GMR}12 (GMR-GAL4), P{w[+mW.hs]=GawB}elav[c155] (elav-GAL4), P{w+mW.hs=GawB}c739 (c739-GAL4), w^1118^;P{w+mW.hs=GawB}c305a (c305a-GAL4), w^1118^;P{w+mW.hs=GawB}1471* (*1471-GAL4*), w[1118]; P{y[+t7.7] w[+mC]=GMR16A06-GAL4}attP2 (*R16A06-GAL4*) and *P{w[+mW.hs]=GawB}3A* (*3A-GAL4*) were obtained from the Bloomington *Drosophila* Stock Center. *w[*]; P{w{+mW.hs]=GawB}NP1131* (*NP1131-GAL4*) and *w[1118]; PBac{EGFP-IV}ank2[KM0104]* (Ank2::EGFP) were obtained from the Kyoto Stock Center. *P{w+mC=tubP-GAL80ts}10* (tubP-GAL80^ts^), p{MEF2-GAL4.247}(*MB247-GAL4*) and *w(CS10)* strains were kindly provided by R. Davis (The Scripps Research Institute, Jupiter, FL). *P{KK106729}VIE-260B* (*UAS-Ank2* RNAi, VDRC ID 107369) and *w^1118^;P{GD12247)v40638 (UAS-Ank2* RNAi, VDRC ID 40638) were obtained from the Vienna *Drosophila* Resource Center. All strains were outcrossed for a minimum of five generations to *w(CS10)* flies. A homozygous line harbouring *w(CS10); P{w+mC=tubP-GAL80_ts_}10* and *P{w+mW.hs=GawB}OK107* (*tubP-GAL80_ts_; OK107-GAL4*) was generated by standard genetic crosses, as was (*elav-GAL4; tubP-GAL80_ts_*), (*c739-GAL4; tubP-GAL80_ts_*), (*tubP-GAL80_ts_, 1471-GAL4*), (*tubP-GAL80_ts_; MB247-GAL4*), (*tubP-GAL80_ts_, c305a-GAL4*) (*tubP-GAL80_ts_; R16A06-GAL4*) and (*tubP-GAL80_ts_, NP1131-GAL4). UAS-DmHDAC4* and *UAS-DmHDAC4 3SA* have been described previously (Main 2021). *UAS-DmHDAC4 ΔAnk* was generated by site directed mutagenesis of *UAS-DmHDAC4* with the following amino acid substitutions: P48A L50A P51A and I53A.

The *UAS-Ank2_190-946_-HA* construct consists of a 2268 bp N-terminal region of *Ank2* containing the ankyrin repeat region (nucleotides 1123 - 3393 of *Ank2*, NCBI reference NM_001274607, which corresponds to amino acids 190 - 946). This construct with a C-terminal 3x HA epitope tag was generated and subcloned into pUASTattB by Genscript (NJ, USA). Transgenic flies were generated by Genetivision (Houston, TX, USA) using the VK22 docking site at 2R(57F5).

### Immunohistochemistry

Whole flies were fixed in PFAT/DMSO (4% paraformaldehyde in 1X PBS +0.1% Triton X-100 +5% DMSO) for one hour then brains were microdissected in 1X PBS. Brains were post-fixed in PFAT/DMSO for 20 mins and blocked in immunobuffer (5% normal goat serum in 1X PBS +0.5% Triton X-100) for three hours prior to incubation with primary antibody of rabbit anti-Ank2-L (1:1000, gift from H. Aberle) (Koch *et al*. 2008), rabbit anti-GFP (Abcam, ab290 1:10,000), mouse anti-nc82 (1:100), mouse anti-Futsch (1:20), mouse anti-Repo (1:20) and mouse anti-Fasciclin II (Fas II, 1:200). Brains were then incubated with secondary antibody (goat anti-mouse Alexa 488 or 555, or goat anti-rabbit Alexa 488 or 555, Sigma Aldrich, 1:500) and mounted with Antifade. The monoclonal antibodies anti-Brp (nc82, developed by E. Buchner), anti-futsch (22C10, developed by S. Benzer and N. Colley), anti-Repo (8D12, developed by C. Goodman), and anti-FasII (1D4, developed by C. Goodman) were obtained from the Developmental Studies Hybridoma Bank developed under the auspices of the NICHD and maintained by The University of Iowa, Department of Biology, Iowa City, IA 52242. For confocal microscopy, images were captured with a Leica TCS SP5 DM6000B Confocal Microscope and images were processed with Leica Application Suite Advanced Fluorescence (LAS AF) software and Image J (NIH). For quantification of dendrite branch length, the total shaft and major branch length inclusive of all six neuronal shafts and their major visible branches were traced using the SNT program in the ImageJ NeuroAnatomy plugin, which allows branching trace plots to be reproduced from the dendritic arborizations (Avery et al., 2017). Total shaft and branch lengths were traced and these measurements were then added together to produce a total sum branch length. Statistical analysis was carried out with the student’s *t*-test with significance set at p<0.05.

### Western Blotting

Whole cell extracts were prepared from 100 snap-frozen heads by homogenizing in RIPA buffer, followed by centrifugation at 13,000 g for 2 minutes at 4°C. Lysates (30 μg) were resolved on 4-20% SDS-PAGE gels (Biorad) and transferred onto nitrocellulose membranes. Membranes were blocked for >1 hour at room temperature in 5% skim milk powder in TBST (50 mM Tris, 150 mM NaCl, 0.05% Tween-20, pH 7.6) then incubated overnight at 4°C in primary antibody, washed 3 × 5 mins in TBST then incubated one hour in secondary anti-mouse, anti-rat or anti-rabbit HRP-conjugated antibodies (GE Life Sciences) as appropriate. Following 3 × 5 min washes in TBST, proteins were detected with Amersham ECL Prime (GE Life Sciences). The following antibodies were used: rabbit anti-GFP (Abcam ab290, 1:10,000); rabbit anti-Myc (Roche, 1:1,000); rat anti-HA (Roche, 1:1,000) and mouse anti-α-tubulin (12G10 clone, DSHB, 1:500).

### Co-immunoprecipitation

Whole cell extracts were prepared as per the western blotting method above. Immunoprecipitation (IP) was performed with the Pierce Classic IP Kit (Thermo Scientific) according to the manufacturer’s instructions. Anti-Myc or anti-HA antibody (1 μL) was incubated overnight with 1 mg of lysate. Following elution in 2x sample buffer, IP samples were processed for SDS-PAGE and western blotting with anti-HA or anti-Myc alongside 30 μg input samples. Anti-α-tubulin (1:500) was used as a loading control.

### RT-qPCR

*elav-GAL4* females were crossed to *UAS-Ank2* RNAi males to generate progeny in which *Ank2* was knocked down in all neurons; and progeny of *elav-GAL4* crossed to *w(CS10)* served as the control. To confirm knockdown, total RNA was extracted from *Drosophila* heads from three independent crosses with the RNeasy Mini kit (Qiagen) according to the manufacturer’s instructions. cDNA was synthesized from 1 μg of total RNA with Transcriptor (Roche) as per the manufacturer’s instructions. RT-qPCR was conducted using SsoFast-EvaGreen (BioRad) reaction master on a Lightcycler II 480 instrument (Roche), following manufacturer’s instructions. The following primers were used: *Ank2for* 5’-GGCCGATATGGCACAAAACC-3’, *Ank2rev* 5’TTCTTTCGACGGTGGTACGG-3’, *EF1a48D*for *5*’-ACTTTGTTCGAATCCGTCGC-3’, *EF1a48D*rev 5’-TACGCTTGTCGATACCACCG-3’. A 5-fold dilution of cDNA from control flies was used as template to prepare a standard curve to confirm efficiency of the PCR reactions. Relative quantification was conducted using 2^−ΔΔCt^ method, normalizing to the housekeeping gene *Ef1α48D* (Livak and Schmittgen 2001). *Ank2* expression was reduced to 0.42 ± 0.12 (mean±standard error) of that of the control, student’s *t*-test t_(12)_=4.74, p<0.001.

### Courtship Suppression Assay

The repeat training courtship suppression assay was used to assess 24-hour long-term courtship memory. This is an experience-dependent assay in which wild-type male flies that have been previously rejected by a mated female will reduce their courtship behavior towards a new mated female. During mating, the male pheromone cVA is transferred to the female, and the presence of this pheromone on the female causes the male to reduce his courtship towards her. Males that have previously experienced rejection will suppress courtship towards another mated female due to an enhanced response to cVA (Ejima *et al*. 2005; Keleman *et al*. 2007) and this form of courtship memory is termed cVA-retrievable memory (Raun *et al*. 2021). The detailed methodology has been described previously (Fitzsimons and Scott 2011; Fitzsimons *et al*. 2013; Freymuth and Fitzsimons 2017). For training, single virgin males (3-5 days post eclosion) of each genotype were placed into individual training chambers. A freshly mated wild-type female was placed with each male to be trained, whereas sham control males were housed alone. Over the seven-hour training period, multiple bouts of courtship were observed in the trained group. The female fly was then aspirated from the training chamber and the males were left in their chambers for the 24 hours, prior to testing. Each trained or sham male fly was then placed into a testing chamber containing a mated wild-type female and was scored for the time spent performing stereotypic courtship behaviors over the ten-minute period. A courtship index (CI) was calculated as the proportion of the ten-minute period spent courting. A mean CI for each group was determined, and from this a memory index (MI) was calculated by the following equation: MI = 1-(CI of each trained fly/mean CI of sham group) (n≥16/group). The MI was measured on a scale of 0 to 1, a score of 0 indicating memory was no different than untrained sham controls. In all experiments, the scorer was blind to the genotype of the flies. For assessment of immediate short-term memory, the training session was reduced to one hour and flies were tested immediately after training. For assessment of learning, the male was placed with a mated female for an hour and the first ten minutes and last ten minutes were scored for courtship behavior. The learning index was calculated as 1-(CI last 10 mins/CI first 10 mins). For statistical analyses, data was arcsine transformed to approximate a normal distribution and one-way ANOVA with post-hoc Tukey’s HSD test was used to assess significance (p<0.05).

### Scanning electron microscopy (SEM)

Flies were anaesthetised using FlyNap (Carolina Biologicals) and fixed in primary modified Karnovsky’s fixative (3% glutaraldehyde, 2% formaldehyde in 0.1 M phosphate buffer, pH 7.2) with Triton X-100 by vacuum infiltration. They were then placed in fresh fixative and incubated at room temperature > 8 hours, followed by 3 × 10 min washes in phosphate buffer (0.1 M, pH 7.2). Dehydration was carried out via a graded ethanol series for ten to fifteen minutes at 25%, 50%, 75%, 95%, and 100% ethanol, followed by a final one-hour incubation in 100% ethanol. The flies were then critical point dried using CO_2_ and 100% ethanol (Polaron E3000 series II drying apparatus). Heads were removed, mounted onto aluminium stubs and sputter coated with gold (Baltex SCD 050 sputter coater). Imaging was performed with a FEI Quanta 200 Environmental Scanning Electron Microscope at an accelerating voltage of 20 kV. To determine the sizes of each eye for comparison between genotypes, ImageJ software was used to draw a line surrounding the eye and calculate the area in arbitrary units. Statistical analysis was performed with one-way ANOVA with post-hoc Tukey’s HSD test with significance set at p<0.05.

To provide a semi-quantitative analysis of SEM images, a scoring system was developed based on observations of phenotypes resulting from overexpression of one or two copies of *UAS-HDAC4* in a previous study (Schwartz *et al*. 2016). No defects: the eye appears wild-type - ommatidia are organised in a regular array with no fusion and mechanosensory bristles are correctly positioned between each ommatidium. Mild: Presence of one of the following phenotypes; between 5 - 10 instances of an abnormal number of interommatidial bristles, mild ommatidial disorganisation or fusion of ommatidia in up to two areas. Moderate: all mild phenotypes were collectively observed or one of the following phenotypes was observed; between 10 – 20 instances of an abnormal number of interommatidial bristles, moderate disorganisation or fusion of ommatidia in up to five areas. Major: all moderate phenotypes were collectively observed or one of the following phenotypes was observed; more than 20 instances of an abnormal number of interommatidial bristles, major disorganisation of the ommatidial array, fusion of ommatidia in up to 10 areas with few large areas of fusion or up to 50 collapsed ommatidia. Severe: all major phenotypes were collectively observed or one of the following phenotypes was observed; severe disorganisation, fusion in more than 10 areas or multiple large patches, more than 50 collapsed ommatidia or severe collapsing of ommatidia resulting in central hole-like cavities. Statistical analysis was assessed with the Fisher’s Exact test with significance set at p<0.05.

## RESULTS

### Characterisation of Ank2 expression in the brain

To date, the expression and localisation pattern of Ank2 has been described in the *Drosophila* neuromuscular junction (Koch *et al*. 2008; Pielage *et al*. 2008) however the expression pattern in the adult brain has not been characterized. Immunohistochemistry on whole mount brains with an antibody that detects Ank2-L (Koch *et al*. 2008) indicated a broad expression profile with high expression in the optic lobes, antennal lobes, mushroom body and axon tracts throughout the brain (Fig 1A,B). Colocalization with the axonal marker Futsch (Hummel *et al*. 2000) confirmed that Ank2-L localizes to axon tracts across the adult brain (Fig 1C). Ank2-L did not co-distribute with the glial marker Reversed Polarity (Repo) (Alfonso and Jones 2002), confirming its specific neuronal expression pattern (Fig 1D). Since the mushroom body is a critical structure for memory (Heisenberg *et al*. 1985; McBride *et al*. 1999), we examined the expression and subcellular distribution of Ank2 in this brain area in more detail. The intrinsic neurons of the mushroom body are the Kenyon cells, which receive input from the olfactory system (Turner *et al*. 2008). The cell bodies of the approximately 2,500 Kenyon cells are clustered in the posterior dorsal region of the brain and extend their dendrites anteriorly into a globular region known as the calyx, which is organized into an array of microglomeruli, each comprising the large synaptic bouton of projection neurons from the antennal lobe surrounded by Kenyon cell dendrites (Leiss *et al*. 2009). Their axons form a bundled fiber termed the pedunculus and project towards the anterior portion of the brain, forming five distinct lobes; the vertical α and α’ lobes and the medial β, β’ and γ lobes (Crittenden *et al*. 1998; Lee and Luo 1999) (see Fig. 3F). We examined the colocalization of Ank2-L with the neuronal cell adhesion molecule Neuroglian (Nrg), which has been shown to interact with Ank2 (Enneking *et al*. 2013; Siegenthaler *et al*. 2015). Nrg is the sole *Drosophila* orthologue of the L1-CAM family of proteins (Bieber *et al*. 1989), which enables axon guidance through the mushroom body. We confirmed that Ank2-L and Nrg codistributed in multiple axon tracts including the axons of the mushroom body, where both were observed in the α, β and γ lobes (Fig 1E). Ank2-L was also concentrated in axon tracts surrounding the calyx of the mushroom body (Fig 1F).

**Figure 1.**
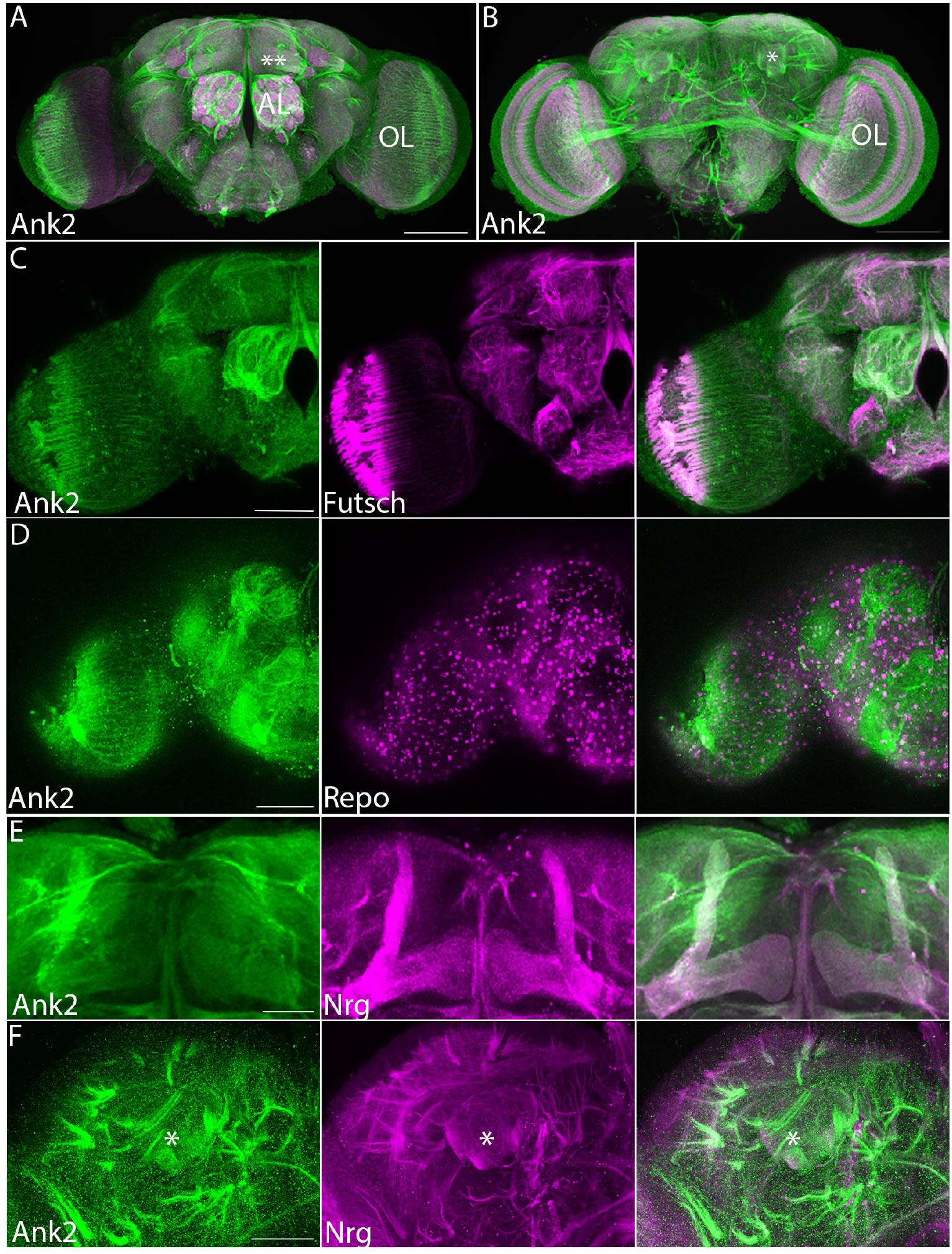
Expression of Ank2 in the adult brain. Immunohistochemistry on whole mount wild-type brains indicates widespread distribution of Ank2-L. A,B Confocal projection of brains labelled with Ank2-L (green) and nc82 (Bruchpilot, magenta) antibodies, to highlight the synaptic neuropil. Images are Z-stacks of 1 μm optical sections. A, Anterior confocal projection. AL, antennal lobe, OL optic lobe, **indicates the γ lobe of the mushroom body. Scale bar = 100 μm B-F. Co-labelled proteins are shown in magenta and labelled in the middle panel. B. Posterior confocal projection, *indicates the calyx of the mushroom body. Scale bar = 100 μm. C. Immunohistochemistry with Ank2-L and 22C10 (Futsch) antibodies showing codistribution in neurons, with widespread localization to axon tracts. Scale bar = 50 μm. D. Ank2-L does not codistribute with pan-glial marker Repo. Scale bar = 50 μm. E. Ank2-L colocalizes with Nrg in the mushroom body lobes. Scale bar = 50 μm. F. Ank2-L also codistributes with Nrg in axon tracts surrounding the calyx (asterisk). Scale bar = 50 μm.

### Ank2 is essential for axon and dendrite morphogenesis

We previously found that overexpression of *HDAC4* impairs morphogenesis of mushroom body axons, with deficits in axon branching and guidance observed as missing α and/or β lobes, misdirected axons as well as the appearance of fused β lobes, resulting from defects in axon termination across the midline (Main 2021). To investigate whether Ank2 is also required for axon morphogenesis, the morphology of the mushroom body in brains with reduced Ank2 was analyzed via detection of Fasciclin II. This cell adhesion molecule is highly expressed in the α, β and γ lobes of the mushroom body (Crittenden *et al*. 1998) and is a commonly used marker to visualise mushroom body lobe architecture (Fig 2A). Pan-neuronal knockdown of *Ank2* with an inverted repeat hairpin that targets all long isoforms of *Ank2* mRNA for degradation resulted in a variety of phenotypic defects of the mushroom body, including thin lobes, missing lobes and guidance abnormalities (Fig 2B-F). GAL4 activity increases at higher temperatures and accordingly we observed more severe defects when the temperature was raised during larval development (Table 1).

**Figure 2.**
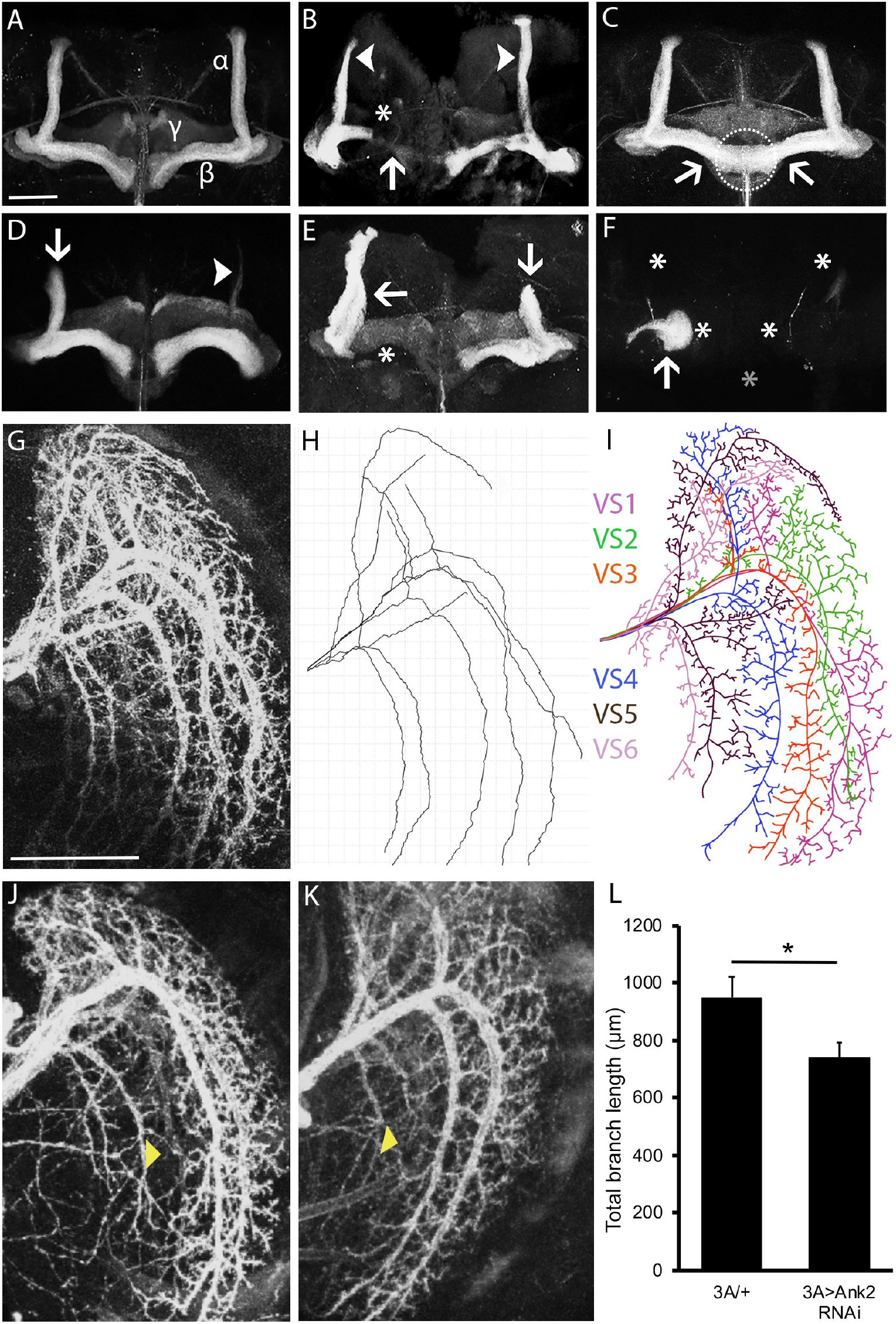
Reduced Ank2 expression disrupts neuronal development. A-F. Immunohistochemistry with anti-FasII on whole mount brains reveals morphological defects of the mushroom body resulting from pan-neuronal expression of *Ank2* RNAi driven by *elav*-GAL4. Knockdown was confirmed by RT-qPCR (as described in the methods). All images are frontal confocal projections through the mushroom body. Scale bar = 50 μm. A. Wild-type mushroom body stained with anti-Fas II to highlight the α, β and γ lobes. B. Thin α lobes (arrowheads), a prematurely terminated β lobe (arrow) and a missing γ lobe (asterisk). C. β lobes (arrows) have crossed the midline and appear fused (circle). D. A thin (arrowhead) and a prematurely terminated α lobe (arrow). E. Misoriented β lobe (arrow) which projects anteriorly rather than its usual medial orientation (asterisk). A prematurely terminated α lobe is also present (arrow). F. None of the lobes have elongated (asterisks) and axon stalling is observed whereby the axons grow in a ball-like structure (arrow). Grey asterisk indicates the midline. G-J. Immunohistochemistry on whole mount brains with anti-GFP driven by *3A-GAL4* to detect Lifeact in LPTCs whole mount brains. All images are confocal projections through the optic lobe of the brain. G. The dendritic arbor of the six neurons comprising the vertical system of LPTCs in a wild-type brain is visualised with anti-GFP. Scale bar = 50 μm. H. Dendritic trace generated using SNT Tracer (Image J). I. Cartoon trace of the confocal micrograph showing the dendritic branching of each of the vertical system neurons. J,K. Knockdown of *Ank2* results in defects in dendritic branching (arrow heads). L. Total dendrite branch length was reduced by knockdown of *Ank2* (student’s *t*-test t_(36)_=2.27, p<0.05).

**Table 1.** Frequency of mushroom body phenotypes resulting from knock down of *Ank2*. The percentage of brains displaying each phenotype was calculated from the total number of brains analyzed for each genotype (n) at 22, 25 and 30°C. Statistical analysis was performed with Fisher’s Exact Test. Knockdown of *Ank2* resulted in significantly more brains with thin lobes (p=0.0032), elongation and guidance deficits (p=0.0032), or absent α and/or β lobes (p=0.0014), than controls at 25°C. The more severe phenotypes of completely absent α and/or β lobes were more pronounced at higher temperatures (27°C compared to 22°C, p=0.0015).

| Genotype<br>Temperature | *elav>w(CS10)*<br>25°C | *elav> Ank2 RNAi*<br>22°C | *elav> Ank2 RNAi*<br>25°C | *elav> Ank2 RNAi*<br>27°C |
| --- | --- | --- | --- | --- |
| *n* | 20 | 30 | 30 | 16 |
| Thin $\alpha/\beta$ or $\gamma$ lobes | 0% | 37% | 23% | 62% |
| Fused $\beta$ lobes | 0% | 13% | 10% | 0% |
| Outgrowth or guidance defect | 0% | 37% | 23% | 50% |
| Absent $\alpha/\beta$ or $\gamma$ lobes | 0% | 6% | 40% | 50% |

During *Drosophila* embryonic and larval stages, *Ank2* mutants exhibit reduced dendritic branching, and in *Drosophila* dopaminergic neurons, knockdown of *Ank2* results in decreased dendritic branching points, leading to a reduced total branch length and a lack of branching complexity (Avery *et al*. 2017). To that end, we investigated whether Ank2 is required for dendrite morphogenesis in the adult *Drosophila* brain. As branching and elongation of Kenyon cell dendrites is difficult to visualize, we instead examined lobular plate tangential cells (LPTCs) of the visual system. The LPTCs are a group of six interneurons in the optic lobe that provide an ideal model system for investigating dendrite growth and branching as they display stereotypical dendritic arborization (Leiss *et al*. 2009). Individual dendrites are easily visualised via expression of Lifeact, a GFP-fused F-actin binding peptide (Riedl *et al*. 2008) with the *3A-GAL4* driver (Scott *et al*. 2002) (Figure 2G), and branch length can be traced and quantified (Figure 2H, I). The characteristic arborization pattern of the six neurons was disrupted by expression of *Ank2* RNAi with severely reduced dendritic projections (Figure 2J, K) leading to reduced total branch length (Figure 2L). These data suggest that wild-type levels of Ank2 are required for both axon branching, guidance and elongation was well as normal dendritic branching and arborization.

### LTM requires Ank2 expression in the γ lobe of the mushroom body

We next assessed whether Ank2 was required for memory formation with the repeat training courtship suppression assay. This test evaluates the memory of a male following exposure to an unreceptive mated female. Following this failure of mating, a male suppresses his courtship activity towards mated females to which he is subsequently presented. After seven hours of training, males form a stable long-term memory that lasts for at least 24 hours (Keleman *et al*. 2007; Fitzsimons and Scott 2011; Fitzsimons *et al*. 2013). After this time, each male is placed with a new freshly mated (unreceptive) female and a courtship index is calculated by dividing the amount of time each male spends courting by the total duration of the observation period. A memory index is calculated by comparing the time a trained male spends courting to that of a sham male. A score of zero indicates that memory is impaired and no different from untrained sham controls, whereas a higher memory index indicates intact memory. This form of courtship memory has been recently described as “cVA-retrievable memory” to differentiate from the associative memory formed when virgin females are used for testing, which uses different circuitry for memory retrieval (Raun *et al*. 2021). Learning and immediate short-term memory were unaffected by pan-neuronal knockdown of *Ank2* (Fig 3A,B). Long-term courtship memory is dependent on an intact mushroom body, therefore unsurprisingly, pan-neuronal knockdown of *Ank2* during development resulted in a significant and severe loss of LTM formation compared to control genotypes (Fig 3C). This was not due to an effect on courtship behavior as sham males of each genotype all spent approximately the same percentage of time courting (87 to 89%, Figure 3D).

**Figure 3.**
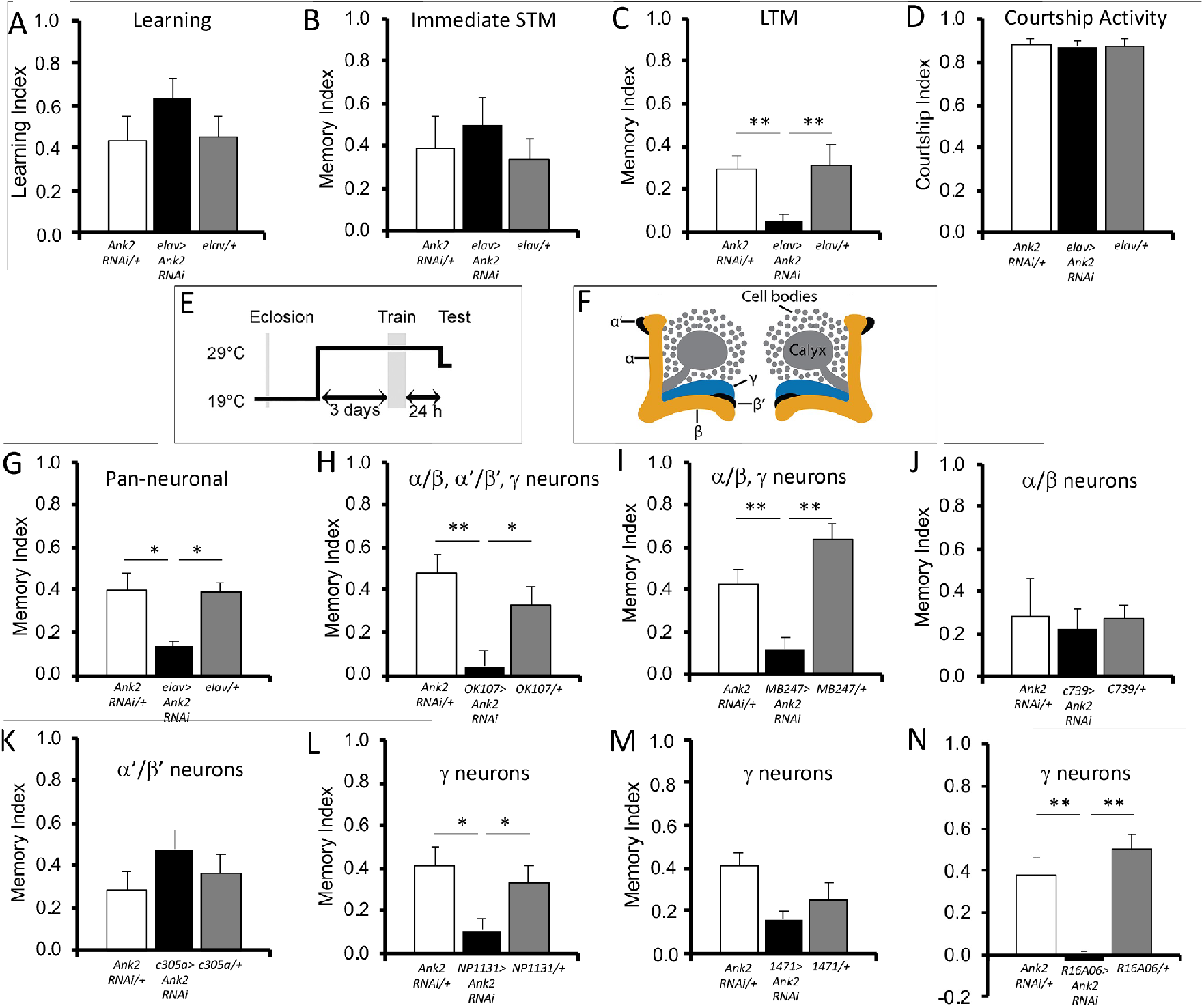
Ank2 is required for long-term memory. Learning and memory were assessed with the courtship suppression assay. The controls included in each assay are the GAL4 driver (plus tubP-GAL80ts where indicated) crossed to *CS*, and UAS-A*nk2* crossed to *CS*, such that the progeny are heterozygous for either the driver or the RNAi. A-D. *elav-GAL4* and UAS-*Ank2* RNAi flies were crossed to achieve panneuronal knockdown of *Ank2* in progeny. A. Learning was unaffected knockdown of *Ank2* (ANOVA, F_(2,47)_=0.002, p=0.252). B. Immediate memory was also unaffected (ANOVA, F_(2,45)_=0.044, p=0.819). C. *Ank2* knockdown impaired long-term memory (ANOVA, F_(2,60)_=7.31, p<0.001; *post-hoc* Tukey’s HSD, **p<0.01). D. Courtship activity was not impaired by pan-neuronal knockdown of *Ank2* (ANOVA, F_(2,51)_=0.14, p=0.870). E-K. Knockdown of *Ank2* in the adult mushroom body impairs LTM. *Ank2* was knocked down in specific regions of the brain by crossing *Ank2* RNAi to the indicated driver line and *tubP-Gal80ts*. E. Schematic diagram depicting the induction of expression in the adult mushroom body. Expression was restricted to the adult brain by raising flies at raised at 19°C, at which temperature GAL80 represses GAL4. After eclosion, when flies were 3-5 days old, the temperature was raised to 30°C for 72 hours, after which training commenced. At this temperature GAL80 is inactivated, allowing GAL4 to induce transgene expression. Twenty-four hours after training, the flies were equilibrated to 25°C for one hour prior to testing. F. Schematic diagram labelling the lobes of the mushroom body in which *Ank2* was knocked down. G. Pan-neuronal knockdown of *Ank2* in the adult brain impairs long term memory (ANOVA, F_(2,54)_=0.317, p<0.01; *post-hoc* Tukey’s HSD, *p<0.05). H. Similarly, memory is also impaired when knockdown of *Ank2* is restricted to the mushroom body (ANOVA, F_(2,52)_=0.922, p<0.001; *post-hoc* Tukey’s HSD, **p<0.01, *p<0.05). I. When knockdown is restricted to the α/β and g neurons of the mushroom body, long-term memory is still disrupted (ANOVA, F_(2,51)_=0.923, p<0.0001; *post-hoc* Tukey’s HSD, **p<0.01). J. Reduction of *Ank2* in just the α/β neurons has no significant effect on long-term memory (ANOVA, F_(2,41)_=0.025, p=0.819). K. There is also no impairment when *Ank2* is reduced in the α’/β’ neurons (ANOVA, F_(2,51)_=0.122 p=0.372). L. *Ank2* is required in the g lobes, as knockdown with *NP1131-GAL4* impairs LTM (ANOVA, F_(2,46)_=0.312, p<0.01; *post-hoc* Tukey’s HSD, *p<0.05). M. The weaker g lobe driver *1471-GAL4* reduced LTM, however this was not quite significant (ANOVA, F_(2,59)_=0.210, p=0.056). Knockdown of *Ank2* with the stronger g lobe driver *R16A06-GAL4* did impair LTM significantly (ANOVA, F_(2,33)_=18.57, p<0.0001, post-hoc Tukey’s HSD, **p<0.01).

To avoid the developmental deficits resulting from decreased *Ank2* expression and allow assessment of the role of Ank2 specifically in adult memory processes, knockdown of *Ank2* was restricted to the mature brain with GAL80ts, a temperature sensitive inhibitor of GAL4 activity (McGuire *et al*. 2004) (Fig 3E). Flies were raised at the permissive temperature of 19°C at which GAL80ts is active. Seventy-two hours after eclosion, male flies from the F1 progeny were collected individually and transferred to 30°C to inactivate GAL80ts and thus induce pan-neuronal RNAi expression. After three days, males were tested in the courtship suppression assay. Adult-specific knockdown of *Ank2* in all neurons resulted in impairment of LTM formation (Fig 3G) and when knockdown of *Ank2* was restricted to the adult mushroom body (Fig 3F) with *OK107-GAL4*, this impairment remained (Fig 3H). The three Kenyon cell subtypes are structurally distinct with individually identifiable transcriptomes (Croset *et al*. 2018) and distinct roles in learning and memory (Joiner and Griffith 1999; Keleman *et al*. 2007), thus we next investigated whether there was a differential requirement for Ank2 in specific mushroom body subtypes by restricting expression of *Ank2* RNAi to each subtype individually (Fig 3F). Expression with the α/β and γ neuron driver *MB247-GAL4* abolished LTM formation (Fig 3I). Knockdown in the α/β neurons or α’/β’ neurons did not significantly alter LTM (Fig 3J,K) whereas knockdown in γ neurons with *NP1131-GAL4* prevented LTM formation (Fig 3L). The γ neuron driver *1471-GAL4* did not quite impair memory to significant levels (Fig 3M), however this is a much weaker than *NP1131-GAL4* (Aso *et al*. 2009). We therefore tested an additional γ neuron driver *R16A06-GAL4* which drives very strong expression in the gamma lobe, but minimal expression elsewhere in the brain (Jenett *et al*. 2012), and employed a second independent *Ank2* RNAi line that also targets all long forms of *Ank2* mRNA, which together resulted in a significant reduction in LTM (Fig 3N).

Taken together these data show that Ank2-L is required for normal mushroom body development and in the adult brain, wild-type levels of Ank2-L are required in the γ lobes for normal LTM formation. This is strikingly similar to the phenotypes resulting from manipulation of HDAC4 expression in that it is also required in the γ lobe for LTM but not STM (Fitzsimons *et al*. 2013).

### HDAC4 does not physically interact with Ank2 in *Drosophila* neurons

The N-terminal region of HDAC4 contains an ankyrin repeat binding domain consisting of a PxLPxI/L motif (Fig 4A), which in mammalian cells binds the ankyrin repeat region of ANKRA2 and RFXANK (Wang *et al*. 2005; Xu *et al*. 2012; Nie *et al*. 2015b). This motif is conserved in *Drosophila* HDAC4, and the ankyrin repeat of RFXANK and ANKRA2 both share 52% amino acid similarity with that of Ank2, thus it is possible that Ank2 and HDAC4 could interact physically through the ankyrin repeat of Ank2. To test for a physical interaction, we generated flies that co-express the ankyrin repeat-containing domain of Ank2 with a C-terminal HA tag (UAS-Ank2_190-946_-HA) and Myc-tagged *Drosophila* HDAC4 in the brain. Coimmunoprecipitation on head lysates revealed no detectable interaction of HDAC4 with Ank2 via pulldown with either anti-HA or anti-Myc (Fig 4B,C), therefore while *Ank2* and *HDAC4* interact genetically, they do not appear to physically interact at a detectable level in the brain. An obvious explanation for the genetic interaction between the two would be that HDAC4 regulates expression of *Ank2*, however previous RNA-seq data in which we expressed DmHDAC4 or a nuclear-restricted mutant of human HDAC4 did not show a significant change in transcription of *Ank2* (Schwartz *et al*. 2016; Main 2021). We also confirmed that increased HDAC4 expression does not alter the level of Ank2 protein (Fig 4D,E).

**Figure 4.**
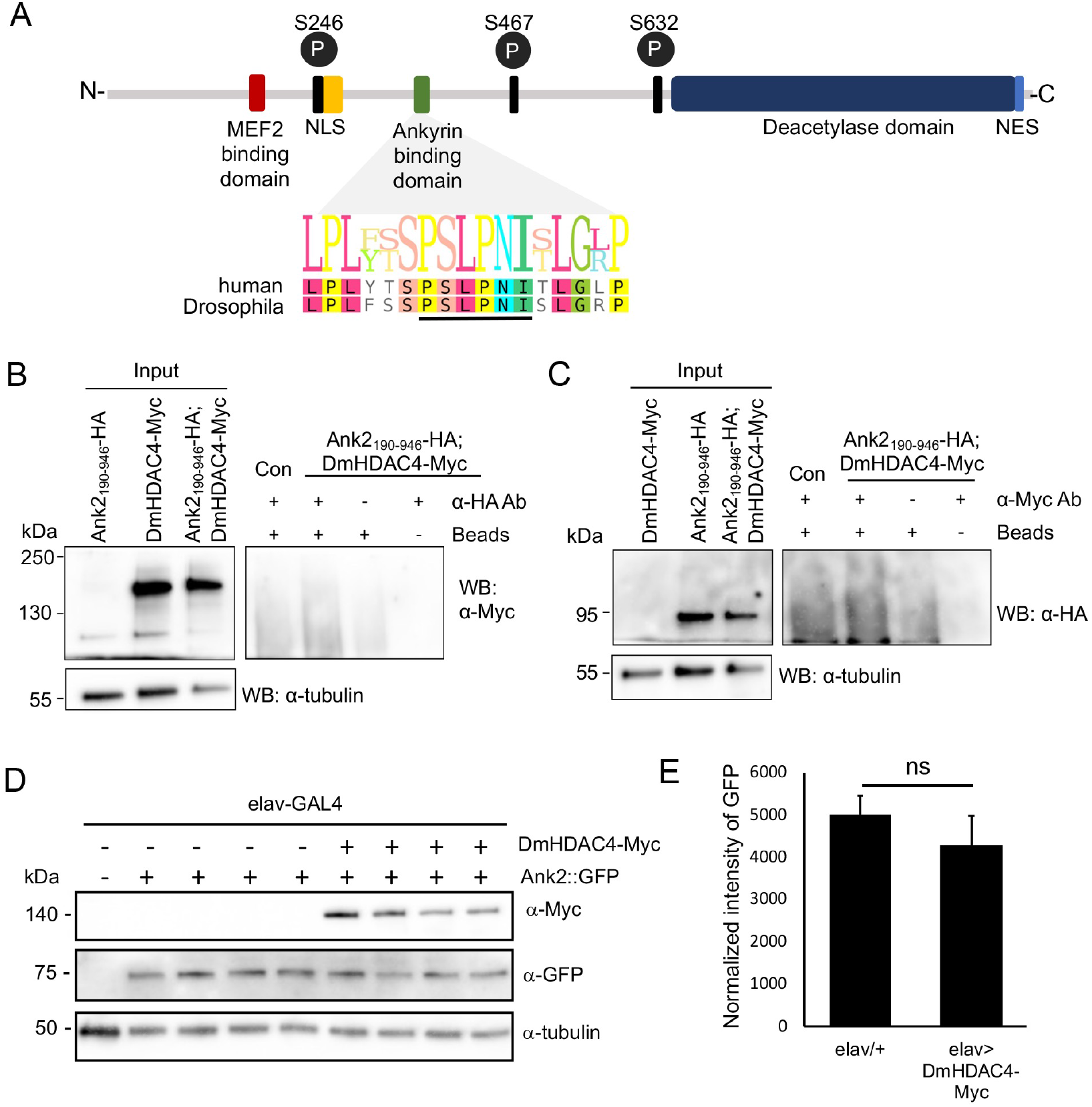
Ank2 does not bind HDAC4 and nor is its expression regulated by HDAC4. A. Domain structure of human HDAC4 showing binding sites conserved between *Drosophila* and human HDAC4. The amino acid sequence of the region containing the RFXANK/ANKRA2 binding site in human HDAC4 is shown, with the corresponding amino acid sequence in *Drosophila* HDAC4. NLS, nuclear localisation sequence, NES, nuclear export sequence. Ps circled in black are serine residues that when phosphorylated provide binding sites for 14-3-3 mediated nuclear export. B,C. Co-immunoprecipitation of Ank2190-946-HA and DmHDAC4-Myc from whole cell lysates of fly heads with either anti-Myc or anti-HA. Both blots were probed with anti-tubulin as a loading control. Input samples = 30 μg. B. Following immunoprecipitation with anti-HA, DmHDAC4-Myc was not detected upon probing with anti-Myc. Inputs include Ank2190-946-HA, which is not detected by Myc, DmHDAC4-Myc, and Ank2190-946-HA; DmHDAC4-Myc to confirm expression. C. In the reciprocal experiment, flies expressing Ank2190-946-HA; DmHDAC4-Myc were subjected to IP with anti-Myc, however Ank2190-946-HA was not detected upon probing with anti-HA. D. *elav-GAL4; Ank2::GFP flies* were crossed to *w(CS)10* and *UAS-DmHDAC4-Myc* and whole head lysates of progeny were generated for western blotting. Samples were processed from four independent crosses. Blots were probed with anti-Myc to verify expression of *DmHDAC4-Myc* and anti-GFP to determine whether the amount of Ank2::GFP normalized to tubulin is altered in the presence of DmHDAC4. E. There was no significant change in the level of Ank2::GFP on expression of *DmHDAC4-Myc*.

### Nuclear HDAC4 mediates the genetic interaction with Ank2

We previously showed that knockdown of *Ank2* in the eye resulted in a mild developmental impairment that was significantly enhanced when combined with *HDAC4* overexpression, indicating the two genes interact in the same molecular pathway (Schwartz *et al*. 2016). The HDAC4-induced impairments in eye development are largely a consequence of nuclear accumulation of HDAC4, as expression of a mutant variant of human HDAC4 that is sequestered in the nucleus resulted in a more severe phenotype than overexpression of wild-type human *HDAC4*, whereas the phenotype resulting from expression of a cytoplasm-restricted mutant was mild (Main 2021). To that end, we next investigated whether the genetic interaction between *Ank2* and *HDAC4* in the eye is also dependent on the nuclear presence of *Drosophila* HDAC4. In addition, we sought to determine whether the genetic interaction is dependent on the putative ankyrin repeat-binding motif region of HDAC4. If not, this would provide further confirmation that the genetic interaction between *HDAC4* and *Ank2* is not through direct physical binding.

We first confirmed our previous observations by co-expressing *DmHDAC4* and *Ank2* RNAi under the control of *GMR-GAL4*, which drives expression in post-mitotic cells posterior to the morphogenetic furrow (Freeman 1996), and examining the phenotype of adult eyes. We also raised the temperature to 27°C to increase GAL4 activity and thus increase the degree of knockdown in order to examine the resulting phenotypic defects in eye development in more detail. Control flies (*GMR-GAL4* crossed to the background *w[CS10]* strain) showed predominantly normal ommatidial alignment and no evidence of fusion (Fig 5A), whereas eyes with reduced *Ank2* expression had fused, collapsed and misaligned ommatidia lacking some interommatidial bristles (Fig 5B). Overexpression of *DmHDAC4* also resulted in missing/disorganised bristles and misaligned and fused ommatidia (Fig 5C). Combined expression of *DmHDAC4* and *Ank2* RNAi resulted in a more severe rough eye phenotype consisting of major areas of ommatidial fusion, severe misalignment with hole-like cavities within the ommatidia, and missing/disorganised bristles (Fig 5D). The severity of phenotypes was scored (Table S1) and the percentage of eyes displaying the most severe phenotype was significantly higher when *DmHDAC4* and *Ank2* RNAi were combined (Fig 5I). In addition to the severe eye phenotypes observed above, the eyes were also significantly physically smaller than those expressing either *Ank2* RNAi or *DmHDAC4* individually (Fig 5J). Together these synergistic phenotypes are consistent with our previous findings (Schwartz *et al*. 2016) and provide further confirmation that *Ank2* and *HDAC4* genetically interact together to influence eye development.

**Figure 5.**
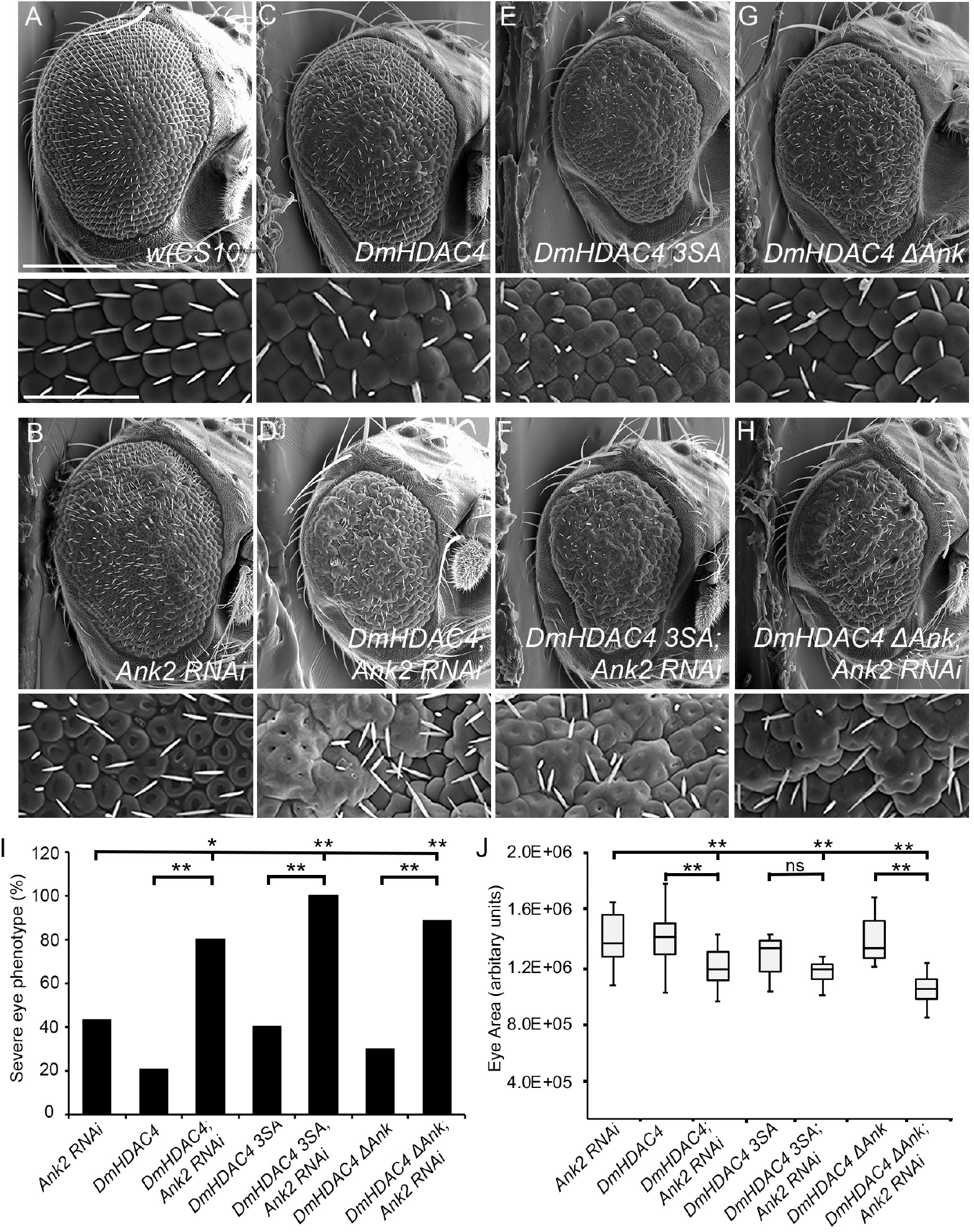
*Ank2* interacts genetically with *HDAC4* in the eye. A-H. Scanning electron micrographs of *Drosophila* eyes expressing *Ank2* RNAi and/or *HDAC4* variants. The genotypes indicated in each panel were generated by crossing *GMR-GAL4* females to males carrying each *UAS-HDAC4* construct or *Ank2* RNAi to the *w(CS10)* control. Top panel: Scale bar = 200 μm. Bottom panel: Scale bar = 100 μm. I. The percentage of eyes displaying severe phenotypes are shown for each genotype. Severe phenotypes were scored as severe disorganisation with fused ommatidia in more than ten areas, and/or more than 50 collapsed ommatidia with crevices or cavities. *p < 0.05, **p < 0.01 following one-tailed Fisher’ s exact test. p-values: *Ank2* RNAi:*DmHDAC4; Ank2* RNAi = 0.0247, *Ank2* RNAi:*DmHDAC4 3SA; Ank2* RNAi = 0.0006, *Ank2* RNAi:*DmHDAC4 ΔAnk; Ank2* RNAi = 0.0028, *DmHDAC4: DmHDAC4; Ank2* RNAi = 0.0004, *DmHDAC4 3SA:DmHDAC4 3SA; Ank2* RNAi = 0.0005, *DmHDAC4 ΔAnk:DmHDAC4 ΔAnk2; Ank2* RNAi = 0.0002. J. Eye sizes were quantified by tracing a line around each eye and calculating the area in arbitrary units using ImageJ software and plotting on Box and whisker plots to show the variation in eye size for each genotype. **p < 0.01 following one-way ANOVA and post-hoc Tukey’s HSD test for significance. p-values: *w(cs10):Ank2* RNAi = 0.007, *w(cs10):DmHDAC4* = 0.009, *w(cs10):DmHDAC4; Ank2* RNAi = 0.001, *Ank2* RNAi:*DmHDAC4; Ank2* RNAi = 0.001, *DmHDAC4:DmHDAC4*; *Ank2* = 0.001, *w(cs10)*:*DmHDAC4 3A* = 0.001, *w(cs10):DmHDAC4 3A; Ank2* RNAi = 0.001, *DmHDAC4 3A:DmHDAC4 3A; Ank2* RNAi = 0.285, *w(cs10):DmHDAC4 ΔAnk* = 0.001, *w(cs10)*:*DmHDAC4 ΔAnk*; *Ank2* RNAi = 0.001, *DmHDAC4 ΔAnk:DmHDAC4 ΔAnk; Ank2* RNAi = 0.001. n = number of eyes per sample. The number of eyes analyzed per genotype is indicated in Table S1.

To further investigate the mechanism of this interaction, we examined whether it was dependent on the nuclear activity of HDAC4. A nuclear-restricted mutant of *HDAC4* (*DmHDAC4 3SA*) resulted in a more severe phenotype than wild-type HDAC4, with increased disorganisation and fusion of ommatidia and a reduced number of bristles (Fig 5E), which we have previously observed (Main 2021). *DmHDAC4 3SA* interacted synergistically with *Ank2* (Fig 5F), confirming that nuclear activity of HDAC4 is required for the genetic interaction. To address whether the putative ankyrin binding motif in *Drosophila* HDAC4 is required we substituted residues that have been shown to be important for this interaction in mammals; P48 L50 P51 and I53 of the PxLPxI/L motif (Fig 4A) to alanines to create the *DmHDAC4 ΔAnk mutant*. Expression of this mutant also resulted in a moderate rough eye phenotype consisting of ommatidial fusion and misalignment (Figure 5G), indicating that the presence of this motif is not required for the *HDAC4* overexpression-induced eye defects. This mutant still interacted genetically with *Ank2*, resulting in a significantly more severe rough eye phenotype and reduced eye size when expressed in combination with *Ank2* (Fig 5H), therefore this interaction does not depend on the presence of the putative ankyrin-binding motif of HDAC4, which is consistent with the lack of physical interaction with Ank2.

## DISCUSSION

Here, we provide the first characterization of the expression pattern of Ank2 in the adult *Drosophila* brain and demonstrate that Ank2 is critical for normal development of the mushroom body and for formation of long-term memories.

Ank2 expression was observed throughout the brain and localized predominantly to axon tracts, including the lobes of the mushroom body where it colocalized with Nrg. As the mushroom body is a critical structure for memory formation (Heisenberg *et al*. 1985; McBride *et al*. 1999), and we previously showed that HDAC4 is required in the mushroom body for normal memory formation, and that overexpression of HDAC4 impairs mushroom body development (Fitzsimons *et al*. 2013; Main 2021), we focused our attention on examining the role of Ank2 in both developmental and post-developmental processes in this brain region. Pan-neuronal knockdown of *Ank2* in the developing mushroom body resulted in deficits in axon guidance and elongation. Ank2 has been demonstrated to interact with Nrg through a FIGQY motif within the intracellular domain of Nrg (Enneking *et al*. 2013). Nrg is required for normal mushroom body lobe development, whereby the extracellular domain of Nrg mediates cellular adhesion between axons of different mushroom body subtypes for guidance into the pedunculus and lobes. This interaction relies on the presence of Ank2 in either the ingrowing or substrate axon neurons where it is proposed to stabilise the transaxonal Nrg complex (Siegenthaler *et al*. 2015). A hypomorphic *Nrg* mutant displayed deficits in axon growth and guidance in the mushroom body (Siegenthaler *et al*. 2015), which are very similar to those we observed on knockdown of *Ank2*. In addition, Nrg also interacts with a second cytoskeletal adapter protein Moesin (Moe) through a FERM domain in the intracellular domain of L1 CAMs (Dickson *et al*. 2002), creating a ternary complex between Ank2, Nrg and Moe (Siegenthaler *et al*. 2015). Moe is highly expressed in the mushroom body and distributes to the lobes on activation by phosphorylation (Freymuth and Fitzsimons 2017), and likely acts to link the Ank2-Nrg complex to the actin cytoskeleton in mushroom body axons. Interestingly we also previously showed that *HDAC4* interacts genetically with *Moe* (Schwartz *et al*. 2016), and moreover, reduction of *Moe* also shows similar disruption to mushroom body development as with *Ank2* and *Nrg*, with defects in axon elongation and guidance (Siegenthaler *et al*. 2015; Freymuth and Fitzsimons 2017). Together these data support the evidence for a functional relationship between Ank2, Nrg and Moe in mushroom body development.

Pan-neuronal knockdown of *Ank2* also severely impaired 24-hour LTM without affecting courtship behavior, learning, or immediate courtship memory. It was previously found that decreased expression of *Ank2* in the mushroom body with *OK107-GAL4* did not result in learning deficits but caused significant impairment to one-hour STM in the same assay (Iqbal *et al*. 2013). Similarly, pan-neuronal knockdown of *Ank2* with *elav-GAL4* impaired one hour olfactory memory (Higham *et al*. 2019) and mushroom body-specific knockdown with *nSyb-GAL4* impaired three-hour memory when tested in the olfactory conditioning assay (Walkinshaw *et al*. 2015). However, in these studies the drivers are expressed in larvae at which time the mushroom body is developing (Nicolai *et al*. 2003; Ogienko *et al*. 2020; Kobler *et al*. 2021), and an intact mushroom body is required for normal olfactory STM (Heisenberg *et al*. 1985) and associative STM lasting longer than 30 mins (McBride *et al*. 1999). Furthermore, output from γ neurons is required for short-term courtship memory (Keleman *et al*. 2012) and knockdown of *Ank2* in γ neurons during mushroom body development was recently found to result in a shortened axon initial segment in third instar larval brains (Spurrier *et al*. 2019). In light of the severe mushroom body defects we observed on knockdown of *Ank2*, and that Ank2 has been implicated in synapse stability (Hortsch *et al*. 2002; Koch *et al*. 2008), it is unsurprising that memory would be impaired, and from these data it cannot be determined whether Ank2 plays a specific role in memory or whether the defects are a result of impaired morphogenesis of the mushroom body. To that end, in order to dissociate developmental effects from the molecular processes required for LTM in an adult brain, knockdown of *Ank2* was restricted to mature neurons with GAL80. A specific deficit in 24 hour LTM was observed when *Ank2* was knocked down in the adult mushroom body, and subsequent testing of GAL4 drivers that restrict expression to specific mushroom body subtypes revealed that knockdown in just the γ neurons was sufficient to impair memory. We previously showed that overexpression of *HDAC4* in γ neurons also impaired the formation of long-term courtship memory (Schwartz *et al*. 2016) and strikingly, in accordance with the similarity in mushroom body defects, knockdown of *Moe* resulted in the same phenotype as *Ank2* knockdown and *HDAC4* overexpression (Fitzsimons *et al*. 2013; Freymuth and Fitzsimons 2017), suggesting that a functional relationship between these proteins also occurs during LTM formation.

The identification that Ank2 is required in the γ neurons of the mushroom body for normal cVA-retrievable memory is consistent with current models of the circuitry that facilitates this memory (Keleman *et al*. 2012; Raun *et al*. 2021). cVA-retrievable memory involves activation of aSP13 dopaminergic neurons which innervate the γ5 compartment at tip of the γ lobe. This results in increased synaptic transmission from γ neurons to glutamatergic M6 output neurons, which themselves feedback to innervate aSP13 neurons to form a recurrent activation loop (Keleman *et al*. 2012; Zhao *et al*. 2018). Long-term memory requires a later reactivation of aSP13 neurons, which is dependent on sleep (Dag *et al*. 2019). Expression of epigenetic regulators required for cVA-retrievable memory, including Rpd3, HDAC4 and G9a has also been pinpointed to a requirement in only γ neurons. Similarly, the cytoplasmic polyadenylation element-binding protein Orb2 is required for translation of synaptic mRNA during LTM consolidation in the y lobe (Kruttner *et al*. 2015).

Although Ank2 and HDAC4 both localize to axons of the mushroom body, and overexpression of *HDAC4* in the mushroom body also results in impaired axon elongation and guidance defects as well as deficits in LTM formation (Fitzsimons *et al*. 2013; Freymuth and Fitzsimons 2017; Main 2021), we found no evidence of a physical interaction in *Drosophila* neurons. We further investigated the interaction between *HDAC4* and *Ank2* in the eye and observed that a nuclear-restricted mutant of *HDAC4* interacted genetically with *Ank2*, however this was not dependent on the presence of a conserved putative ankyrin-binding motif in HDAC4. Given the striking similarity of the *Ank2* and *Moe* knockdown phenotypes with *HDAC4* overexpression, it is possible that when increased in abundance in Kenyon cell nuclei, HDAC4 may act indirectly to disrupt the normal role of the Ank2, Nrg, and Moe complex. It is not yet known how an HDAC4-mediated signal from the nucleus modulates these or other proteins involved in neuronal morphogenesis and LTM. We previously found that HDAC4 also interacts genetically with several components of the SUMOylation machinery, including the SUMO E2-conjugating enzyme Ubc9, which is also required for long-term courtship memory and interacts genetically with HDAC4 during this process (Schwartz *et al*. 2016). HDAC4 has been proposed to act as an E3 ligase to enhance SUMOylation of target proteins (Gregoire and Yang 2005; Zhao *et al*. 2005). Given that SUMOylation regulates neuronal protein activity (Henley *et al*. 2014) and regulates memory formation (Yang *et al*. 2012; Chen *et al*. 2014; Lee *et al*. 2014; Yu *et al*. 2020), it would be worthwhile to further investigate whether any of the HDAC4-interacting genes are SUMOylated and whether this is impacted by altering the expression level or subcellular distribution of HDAC4.

In our screen for genes that enhanced the *HDAC4*-induced rough eye phenotype (Schwartz *et al*. 2016), the largest group of genes we identified composed of regulators of the actin/spectrin cytoskeleton such as *Trio, NetrinB, Derailed, Ankyrin, Ankyrin2*, and *Moesin*, suggesting that *HDAC4* might regulate memory through interaction with genes involved in remodelling of the actin/spectrin cytoskeleton. This is a phenomenon which is believed to regulate the structural changes including polymerization/depolymerization or alterations of the underlying actin cytoskeleton that underpin learning and memory (Engert and Bonhoeffer 1999; Krucker *et al*. 2000; Lamprecht and LeDoux 2004; Leiss *et al*. 2009; Ojelade *et al*. 2013; Lamprecht 2014). We previously showed that expression of a constitutively active mutant of *Moe* increases the density of F-actin in spine-like structures in LPTC neurons (Freymuth and Fitzsimons 2017). In mammals, AnkG is present at synapses and is required for formation of nanodomain structures in the perisynaptic spine and spine neck of dendrites, the contents of which include adhesion molecules, F-actin and CaMKII, thus facilitating interaction with the actin cytoskeleton for F-actin rearrangements that underlie dynamic alterations in spine morphology. The geometry of the spine in this region increases on induction of long-term potentiation in primary cortical neurons, and reduction of AnkG prevents spine enlargement (Smith *et al*. 2014). It is also notable that inhibition of actin polymerization within the mushroom body of the honeybee enhanced associative olfactory memory (Ganeshina *et al*. 2012). As reduced HDAC4 also impairs LTM, indicating an essential role in memory formation, the impact of both increased and reduced HDAC4 on cytoskeletal rearrangement warrants further investigation.

## Acknowledgements

We thank Hermann Aberle (Heinrich-Heine-Universtität, Düsseldorf, Germany) for the Ank2-L antibody and Max Scott for constructive comments on the manuscript. We also thank the Manawatu Microscopy and Imaging Centre, Massey University for assistance with confocal and SEM.

