## Supplementary material for "Ankyrin2 is required for neuronal morphogenesis and long-term memory and interacts genetically with HDAC4": Table S1

| Phenotype | No defects | Mild | Moderate | Major | Severe | n |
|---|---|---|---|---|---|---|
| *GMR>w(CS10)* | 71% (15) | 29% (6) | 0% | 0% | 0% | 21 |
| *GMR>Ank2* RNAi | 5% (1) | 14% (3) | 14% (3) | 24% (5) | 43% (9) | 21 |
| *GMR>DmHDAC4* | 0% | 11% (2) | 47% (9) | 21% (4) | 21% (4) | 19 |
| *GMR>DmHDAC4;Ank2* RNAi | 0% | 0% | 0% | 20% (4) | 80% (16) | 20 |
| *GMR>DmHDAC4ΔAnk* | 0% | 0% | 50% (10) | 20% (4) | 30% (6) | 20 |
| *GMR>DmHDAC4ΔAnk; Ank2* RNAi | 0% | 0% | 0% | 11% (2) | 89% (17) | 19 |
| *GMR>DmHDAC4 3SA* | 0% | 0% | 10% (2) | 50% (10) | 40% (8) | 20 |
| *GMR>DmHDAC4 3SA;Ank2* RNAi | 0% | 0% | 0% | 0% | 100% | 13 |

**Table S1. Frequency and severity of rough eye phenotype resulting from knockdown of *Ank2* and expression of wild-type and mutant *HDAC4*.** The percentage of eyes displaying each phenotype was calculated from the total number of eyes for each genotype (n).
